# Correlating predicted epigenetic marks with expression data to find interactions between SNPs and genes

**DOI:** 10.1101/2020.02.29.970962

**Authors:** Antoine Despinasse, Yongjin Park, Michael Lapi, Manolis Kellis

**Affiliations:** Computer Science and Artificial Intelligence Laboratory, Massachusetts Institute of Technology, Cambridge, Massachusetts 02139, USA; Sorbonne Universités, Paris, France; Université Côte d’Azur, Nice, France

## Abstract

Despite all the work done, mapping GWAS SNPs in non-coding regions to their target genes remains a challenge. The SNP can be associated with target genes by eQTL analysis. Here we introduce a method to make these eQTLs more robust. Instead of correlating the gene expression with the SNP value like in eQTLs, we correlate it with epigenomic data. This epigenomic data is very expensive and noisy. We therefore predict the epigenomic data from the DNA sequence using the deep learning framework DeepSEA (Zhou and Troyanskaya, 2015).

## INTRODUCTION

Genome wide association studies (GWAS) are a powerful tool to study common, complex diseases. They help find genetic variants associated with a disease. Most efforts to analyze GWAS data are confined to single-nucleotide polymorphisms (SNP) occurring in coding regions of the genome. However, most SNPs found in GWAS are in non-coding regions (Deplancke et al., 2016; Gandal et al., 2016; Ward and Kellis, 2012). This has led to the hypothesis that variant SNPs in non-coding regions cause a change in gene expression rather than in protein function (Tak and Farnham, 2015). The first step in identifying the phenotypic consequences of these variants is to identify their respective gene targets.

Identification of cis-regulatory elements and their gene targets was found to be a difficult task given the dimension of the non-coding genome (ENCODE Project Consortium, 2012). The effects of theses regulatory elements can be held over considerable distances (Chris Cotsapas, 2018), making proximity-based assignments incorrect.

Several approaches have emerged as solutions to the problem: identifying genes with an eQTL driven by a disease risk variant in a locus, and identifying genes affected by regulatory elements driving disease risk (Chris Cotsapas, 2018).

These approaches have limitations. eQTLs are common and therefore do not show causality. They are also very noisy. Close eQTLs are also not independent because of all the linkage disequilibrium (LD).

Epigenomic marks influence gene expression so another approach is to correlate the epigenomic data with the gene expression in order to discover important regulatory regions for each gene. Unfortunately, this epigenomic data is very expensive and noisy. To tackle these two issues, we predict the epigenomic data from the DNA sequence using DeepSEA (Zhou and Troyanskaya, 2015).

DeepSEA (Zhou and Troyanskaya, 2015) is a deep learning-based algorithmic framework for predicting the chromatin effects of sequence alterations with single nucleotide sensitivity. DeepSEA can accurately predict the epigenetic state of a sequence, including transcription factors binding, DNase I sensitivities and histone marks in multiple cell types. The predicted epigenetic state’s only cost is the computation cost, it is more relevant to our use than the measured data because it is more robust and not sensitive to individual measurements perturbations. The model is made available through Kipoi’s API (Avsecetal., 2018).

Our methodology aims to correlate the predicted epigenetic information with expression measurements. Using this, we create a gene network of interaction for transcription factors (TF).

## METHODS

All the code for our method can be found on GitHub (https://github.com/adespi/link_epi_to_expr)

We use the sequence and expression data from the 1000 genome project (Consortium and The 1000 Genomes Project Consortium, 2015; Consortium and GTEx Consortium, 2017). We have both the expression and the sequence solely for 445 because we use the expression from Geuvadis project and take only the European American individuals to reduce confounding factors.

We can analyze the expression by PCA and remove its 5 first components in order to reduce confounding factors (as biggest components usually contain mostly batch effects). We can take the log of the gene expressions to match the biological meaning of the correlation. We can standardize the gene expressions to give them an equal value in the correlation. To fulfill all these goals, we take a version of the expression pre-processed by the Geuvadis team called “GD462.GeneQuantRPKM.50FN.samplename.resk10.txt.gz”.

Due to the high number of genes in humans, we select only a few TFs (transcription factors) to test. We start as a basis with the list from the interaction network between TFs and genes found by Marbach et al. (2016) while working on Tissue-specific regulatory circuits. By using this list, we reduce the amount of data to process and keep only TFs that have a higher probability of being interesting. We have the Geuvadis expression data only for 392 TFs out of the 633 that we found in the paper by Marbach et al. (2016).

For each TF, we extract a 500kb fasta sequence around the gene start from the vcf files, only taking into account the SNPs and not indel, mnp, ref, bnd, or any other alteration from reference sequence. We output only 1 sequence per individual even though they all have two alleles. We take the sequence of the alternate allele in case of heterozygous genotype.

### First round

#### DeepSEA predictions

We give this sequence to the DeepSEA model which runs through the 500kb around the gene start with a window of 1kb and a step of 100bp making 5000 predictions per gene (see Figure 1).

**Figure 1.**
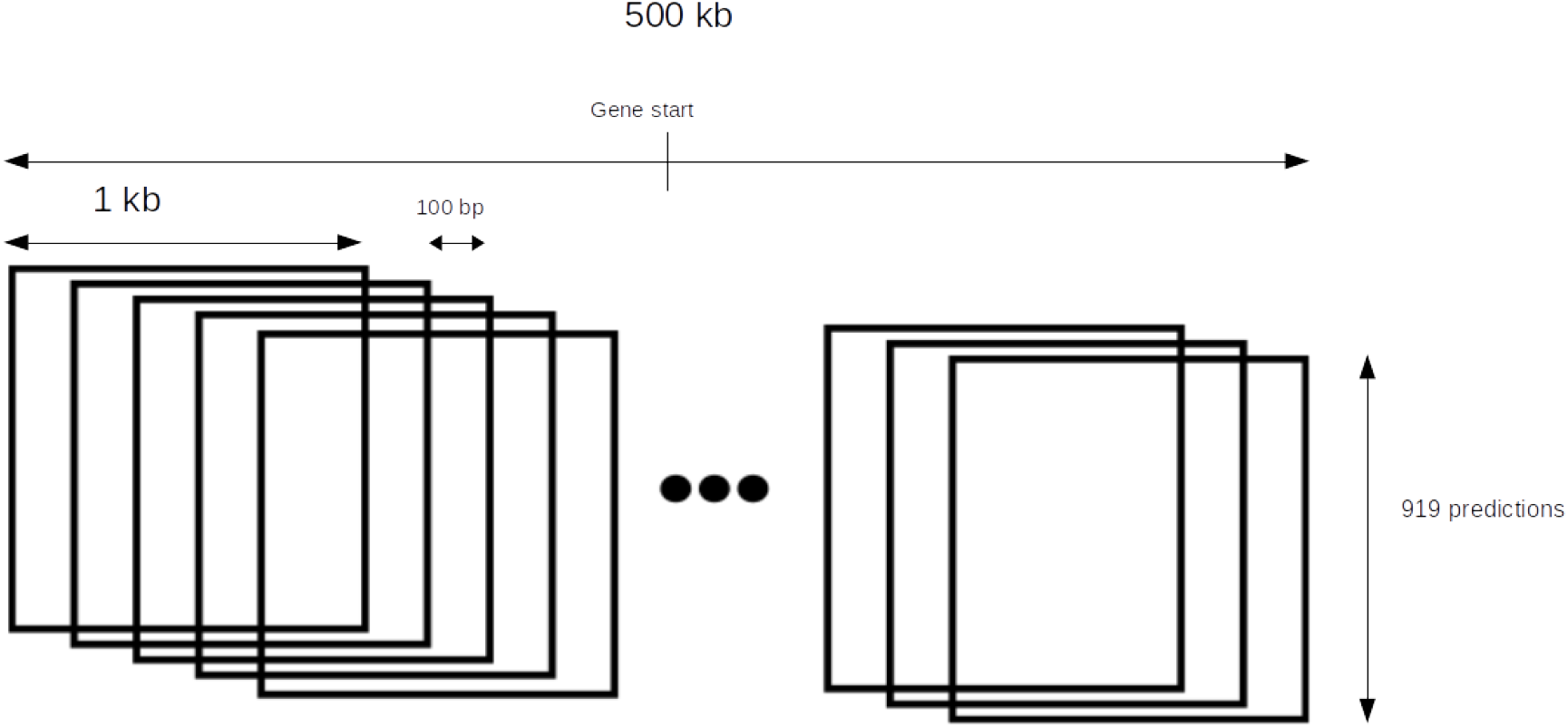
Windows organization around gene start

To compute all these predictions, we started with a CPU computer and the command line interface (CLI) version of Kipoi. The CLI only allows 9 predictions at a time and needs to rebuild the model each time. Building the model took more time than the actual prediction so we moved to the python version of Kipoi, which builds the model once before being able to use the model as many times as needed. This enables a significant jump in speed from 2 iterations/sec to 12 iterations/sec (all the times are in CPU time). A further speed improvement is gained by using GPUs for the predictions enabling 47 iterations/sec. However, we notice that this configuration with the GPU is spending most of its time on data loading and pre-processing because these actions are performed by the CPU. Our current configuration gives predictions for each individual. However, lots of individuals have the same DNA sequence because there are usually few SNPs in a 1000 bp window. The prediction could be calculated only once for different individuals with the same sequence. We tried to implement this unique prediction, but the pre and post processing computation time was higher than the gained time on the prediction. We however don’t calculate positions where there is no SNP at all because that condition is easy to verify. We perform all our time tests on small data-sets and taking a larger data-set gives better results (we suppose because of the optimization from the processors and python for large vectors). Our final model predicts for the 500kb around the gene in 11h of CPU time. On a 16 core machine this corresponds to 40 minutes.

#### Correlations

From the prediction we compute correlations. For this we define:

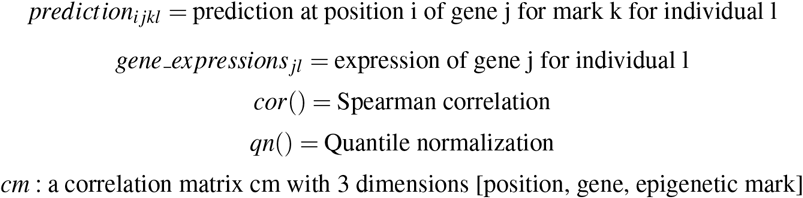

Finally, we calculate cm for each *i, j* and *k* as in equation 1:

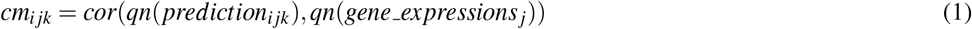

For each position, we find the correlation between its predicted epigenetic marks and the TF’s expression across the individuals (see Equation 1). We tried two languages to calculate the correlation and found that python takes 10 minutes and R 23 minutes to compute all the correlations. We also found that outsiders in the data sometimes give a high false positive correlation. To ensure robustness, the correlations are performed after a quantile normalization using scikit-learn (Pedregosa et al., 2011). As stated in their documentation, this method transforms the features to follow a uniform or a normal distribution. Therefore, for a given feature, this transformation tends to spread out the most frequent values. It also reduces the impact of (marginal) outliers. It may distort linear correlations between variables measured at the same scale but renders variables measured at different scales more directly comparable. In terms of computation time, this increases the correlation CPU time from 10 minutes to 6 hour per 500kb interval (22 minutes on our 16 cores machine).

At this point, we try to find which 1000 bp interval is the most correlated between the delta in expression across the individuals and the delta in DeepSEA prediction. DeepSEA bases it’s prediction on its training data. It may be based on motifs recognition if they are relevant, but can also capture more abstract information, unknown to us.

DeepSEA’s predictions are distributed upon 148 different cell types. We keep all cell type predictions and not only cell types matching with the whole blood expression for several reasons. Firstly, DeepSEA may have missed something in its blood prediction and could have found more motifs or patterns in other cell lines but its results may still relevant for blood. Secondly, a change in whole blood expression can originate from a mutation which has an effect in a tissue. Thirdly, if the expression is changed in blood, it may change in other cell types as well.

As the output number of correlations is very high (5000 *positions* * 919 *marks* * *nbr_of_genes*), we use fdrtool (Strimmer 2008), an R package to compute q-values from the correlations using an empirical model to predict false discovery rate (FDR). The tail area-based FDR is simply a P-value corrected for multiplicity, whereas local FDR is a corresponding probability value. We used the tail area-based FDR in our analysis. The local FDR is more robust but we didn’t have time to rerun the whole process with this FDR. We get q-values for each gene, at each position and for each predicted mark. We keep only the smallest q-value per position and only positions with a *q-value* < 0.05. The correlation for which *q* = 0.05 is 0.20 on average (std of 0.06).

### Second round

After the first round, we have a list of potentially interesting gene positions. For these positions, we compute the correlations between the predictions from DeepSEA and the gene expressions for all the TFs in our list (and not only for the corresponding gene like in the first round). We get a matrix of size (*position* * *TF* * *DeepSEA_prediction).* From these big correlation matrices, we built an interaction network. We consider that two genes are linked if at least one correlation is above the threshold between one gene’s DeepSEA prediction and the other gene’s expression. The threshold for a significant correlation is set at the correlation value 0.2 (correlation for which BIN1 gets a *q – value* < 5 on the first round). We take the interaction network and run a Louvain community detection algorithm (Aynaud, 2011) on it. We group different TFs together (see Results Section).

## RESULTS

### Correlation for position

We look at the best correlations per position to see which regions around the gene are more important in its regulation.

For some genes, we can see some regions standing out on Figure 2a. For BIN1, a region of approximately 50000 bp is clearly more correlated with the gene expression. This region has more impact on the gene’s expression and contains more elements to regulate it.

On the other hand, for other genes like ESRRA, there is no specific region standing out which is more important in the gene regulation (see Figure 2b). On Figure 2c,d we have an example of the best correlation for BIN1. Our method is able to determine the regions near the gene start which are more inclined to be important in the regulation of the gene. We think that these have a high probability of containing an enhancer linked with the promoter of the gene.

**Figure 2.**
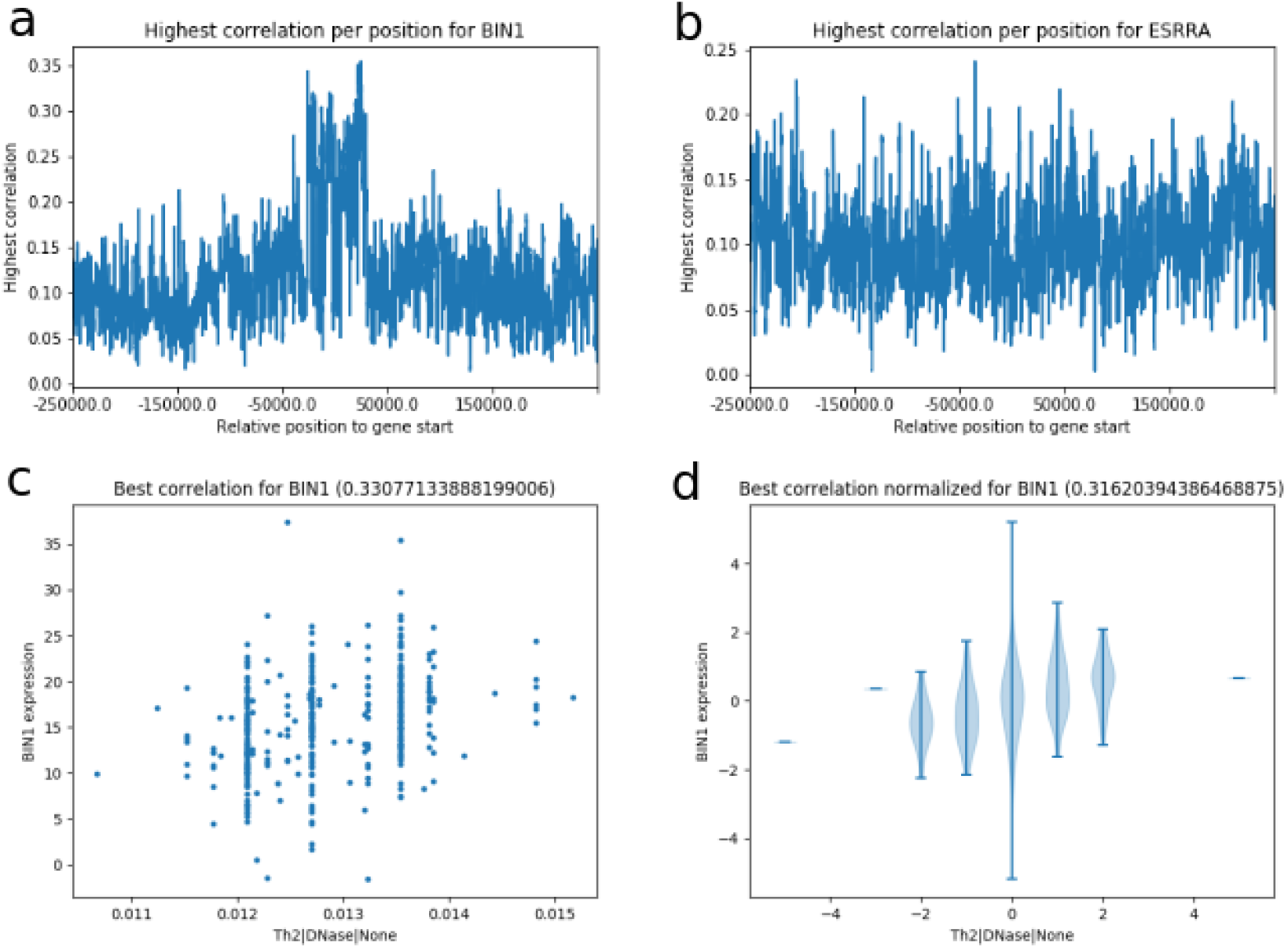
Highest correlation per position and position with best correlation

### Epigenetic marks, cell lines and prediction magnitude

#### Epigenetic marks

We then analyze the significant predictions to see which epigenetic marks are involved and in which cell lines they appear more often. Figure 3a shows the mean correlation depends on the max correlation at each best position per gene. We can see that usually, when one correlation is high for one mark at a given position, all the correlations at this position tend to be high as well. The two variables are highly correlated (Spearman correlation of 0.88) and this shows that there is not one prediction from DeepSEA that is much more significant than the others, but that they more or less all vary in the same way. We did not expect this outcome because we wanted only one (or few) prediction(s) to be positive: the prediction(s) which would tell us which transcription factor’s binding site or epigenetic mark is modified at that precise position. However, the result that we obtain can be explained. Indeed, the DeepSEA predictions come from DNA sequence. If there is only 1 SNP in the 1000 bp window, all the predictions will have only two possible values and the correlation will be exactly the same for all DeepSEA predictions, independently of the range of the epigenetic prediction (because of the normalization of the data necessary for the correlation). A smaller but similar effect can be observed if there are only a few SNP in the 1000 bp window.

**Figure 3.**
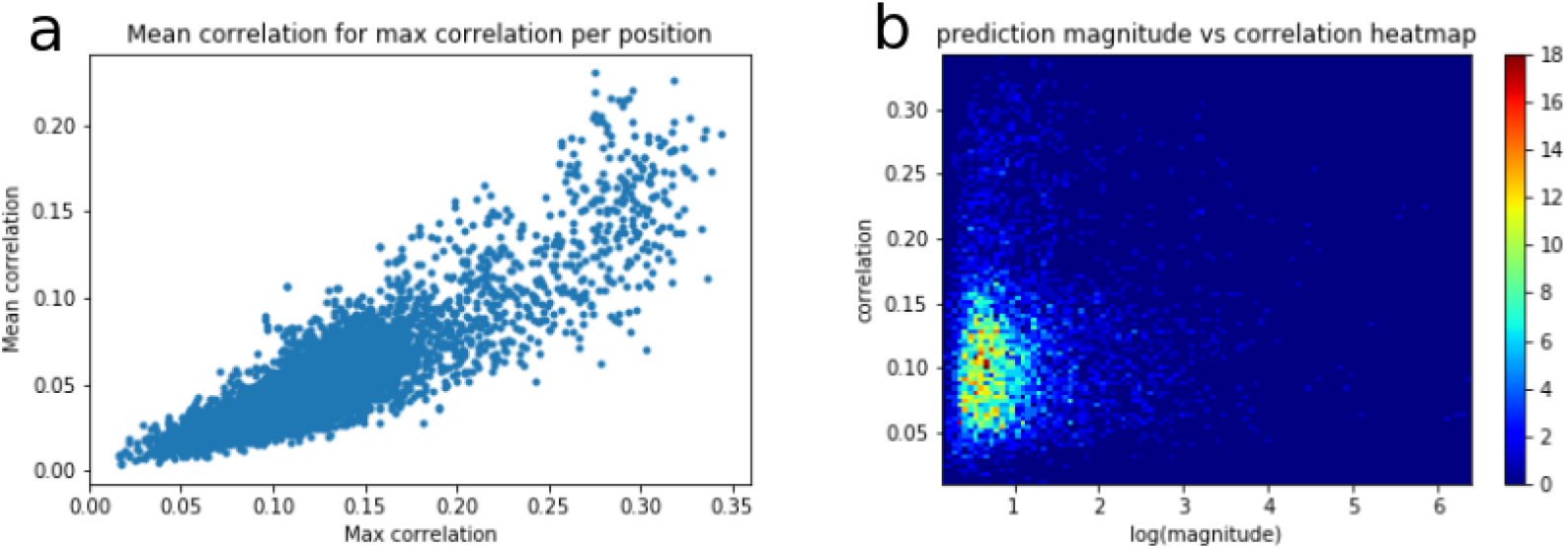
Plot and heat-map for mean correlation, max correlation and prediction magnitude for BIN1

#### Cell lines

The Geuvadis gene expression is from whole blood RNA sequencing. We therefore expected the correlations to be mostly significant with prediction markers from DeepSEA that are on whole blood cell lines. 18.1% of the DeepSEA prediction markers come from whole blood cell lines. However, over all our genes, 17.4% of the significant correlations come from whole blood cell lines.

Similarly, BIN1, is a TF probably involved in AD (Alzheimer’s disease). 4.3% of the DeepSEA prediction markers come from brain cell lines. However, 5.6% of the significant correlations come from brain cell lines.

#### Prediction magnitude

By looking at Figure 3b, we can observe that most of the significant correlations (usually found above the 0. 18 threshold) have a small range of magnitude in the DeepSEA predictions. We interpret that as being a simple eQTL effect, with no added value from DeepSEA. However, we can identify that some significant correlations come from a high magnitude in the DeepSEA predictions. For these data points, we could look at which TF binding site or epigenetic mark have a high log magnitude. This could give us some insight about what is happening inside the cells.

### TF interaction network and gene ontology enrichment

From the method, we obtain the TF interaction network on Figure 4 left. We remove the TFs with no interaction and run a Louvain community detection algorithm (Aynaud, 2011) on it. We group different TFs together on Figure 4 right.

We can see that we obtain 5 clusters of TFs with a graph modularity of 0.13. See clusters here:

1. TBX19, ATF7, HSF1, TGIF2, SOX30, E2F7, HINFP, FOXK1, RFX2
2. PRDM4, YY2, ZKSCAN3, NFE2L1, ATF1, EMX1, RHOXF1, HMX2, MNT, HIC1
3. ATF3, ELK3, BHLHE22, ARX, SOX15, TEF, MYBL2, PPARA, GABPA, IRF5
4. NKX6-3, MIXL1, RARB, GMEB1, RARG, SOX8, BPTF, ATF4, MEF2A, BIN1
5. ZSCAN4, IRF2, ETS2, POU6F1, SIX4, IRF6, PAX8, NR2F6, SOX7, IKZF2, PPARG, ZNF232

**Figure 4.**
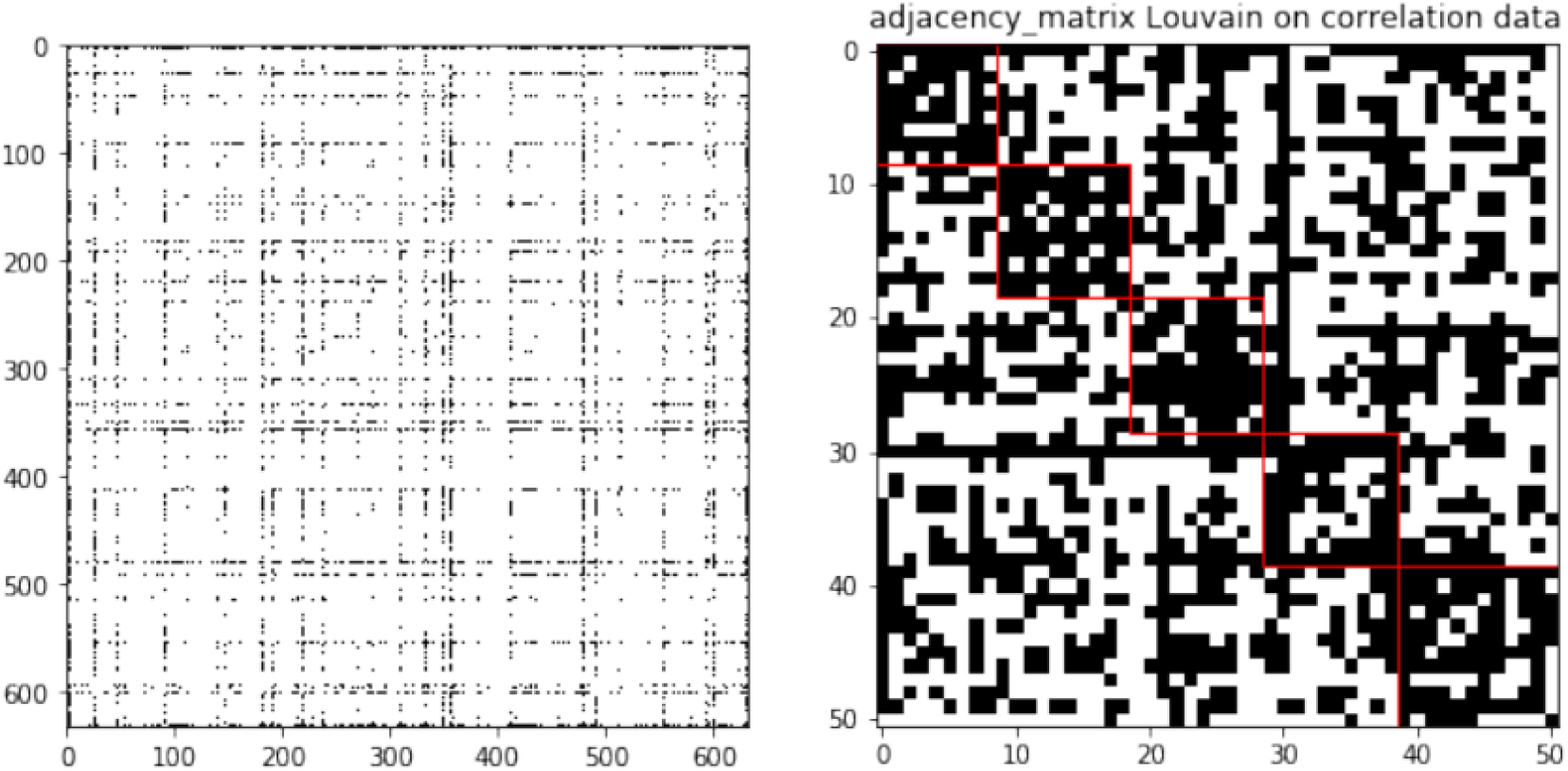
TF interaction network (left) and Louvain communities (right)

Even though the number of genes in each cluster is small (9-12), we perform a gene ontology (GO) enrichment analysis using Panther (Mi et al., 2013) on each of the clusters. We look at which biological processes are affected by the genes in each cluster. See the impacted biological processes in each cluster here:

- Cluster 1
  – regulation of gluconeogenesis by regulation of transcription from RNA polymerase II promoter
  – canonical glycolysis
  – regulation of transcription involved in G1/S transition of mitotic cell cycle
  – negative regulation of transcription by RNA polymerase II
  – chordate embryonic development
  – positive regulation of transcription by RNA polymerase II
- Cluster 2
  – regulation of transcription by RNA polymerase II
  – positive regulation of nucleic acid-templated transcription
  – positive regulation of gene expression
- Cluster 3
  – negative regulation of transcription by RNA polymerase II
  – positive regulation of transcription by RNA polymerase II
- Cluster 4
  – retinoic acid receptor signaling pathway
  – ventricular cardiac muscle cell differentiation
  – glandular epithelial cell development
  – negative regulation of chondrocyte differentiation
  – embryonic hindlimb morphogenesis
  – negative regulation of potassium ion transport
  – growth plate cartilage development
  – embryonic eye morphogenesis
  – regulation of myelination
  – positive regulation of transcription by RNA polymerase II
  – positive regulation of apoptotic process
  – transcription, DNA-templated
  – positive regulation of developmental process
  – negative regulation of transcription, DNA-templated
  – nervous system development
- Cluster 5
  – positive regulation of branching involved in ureteric bud morphogenesis
  – negative regulation of transcription by RNA polymerase II
  – positive regulation of transcription by RNA polymerase II

We see that two clusters contain specific GO enrichment. Cluster 1 mainly contains glucose metabolism and cell replication biological processes. Cluster 4 is composed of embryology and development biological processes. Clusters 2, 3 and 5 mostly contain generic biological processes common to every cell.

## DISCUSSION

Our results tend to show that our method is sensitive to any eQTL effect and captures these in the analysis. There does not seem to be any epigenetic-dependent effect on the correlation score. Our results therefore do not show any direct improvement of our method to eQTL analysis. However, we can focus our attention on genes where there are epigenetic effects involved, even if they do not influence the correlation. There are two ways to do that:

1. Calculate the correlations solely for genes and positions where there is a high difference in DeepSEA prediction between individuals.
2. Calculate all the correlations and focus our interest only on those who have a high difference in DeepSEA prediction between individuals.

Performing this analysis could give us some insight on what is happening inside the cells. This could help us understand which TF binding sites or epigenetic marks are important in the expression of which genes.

